# Expanding the *Egbenema* Horizon: Novel Species Discovery in India and First Complete Genome Assembly of the Genus

**DOI:** 10.1101/2025.03.03.641212

**Authors:** Jayesh M. Bhagat, Ajinkya A. Khilari, Dhanasekaran Shanmugam, Meena Patankar

## Abstract

The recognition of *Egbenema bharatensis*, a novel species of cyanobacteria from the Northern Western Ghats, extends the phylogenetic and ecological understanding of the genus *Egbenema* and highlights terrestrial microbial diversity. Morphological characterization revealed resemblance with other *Egbenema* species but was inconclusive for definitive classification. Initial 16S rRNA sequencing indicated relationship at the genus level, yet sequence similarity less than 98% and observed morphological differences suggested taxonomic novelty. Whole-genome sequencing provided full resolution, with low similarity to known cyanobacterial taxa, and confirmed the isolate as a novel species. Further 16S-ITS region analyses corroborated its novelty and revealed characteristic structural features. Functional genome annotation revealed nitrogen fixation, cobalamin biosynthesis, and stress tolerance pathways, suggesting its metabolic versatility and potential biotechnological applications. The presence of multiple toxin-antitoxin systems and bioenergy-producing genes also reflects its adaptability to environmental fluctuation. This study also presents the first complete genome of the genus *Egbenema*, filling a significant gap in cyanobacterial genomic resources. These findings encourage integrative taxonomic practices and affirm the significance of exploring biodiversity hotspots for uncovering cryptic microbial diversity with ecological and industrial potential.

## Introduction

Cyanobacteria form a diverse and ecologically vital group of photosynthetic microorganisms, playing a fundamental role in nutrient cycling, primary productivity, and nitrogen fixation. Their presence is particularly significant in terrestrial habitats such as soil, tree bark, and rock surfaces, where they exhibit remarkable adaptability to extreme environmental conditions—including desiccation, high UV radiation, and fluctuating temperatures (Geethu & Shamina, 2021). These unique survival strategies not only shape microbial ecology but also position cyanobacteria as valuable resources for biotechnological applications (Zahra et al., 2020).

The Western Ghats of India, a globally recognized biodiversity hotspot, harbor an extraordinary array of flora, fauna, and microorganisms (Ray & Thomas, 2012; Suresh et al., 2012). Despite their ecological richness, only 160 species of cyanobacteria have been recorded in this region—accounting for merely 6% of the global estimate and 57% of the known Asian diversity (Sharathchandra & Sridhar, 2024). Interestingly, many of these species have been documented in polluted environments, underscoring their resilience and adaptability. However, terrestrial cyanobacteria from the Indian subcontinent—particularly those inhabiting soil and tree bark—remain largely unexplored. This is especially true for the Northern Western Ghats, a region with diverse climatic and ecological gradients that presents an exciting opportunity for discovering novel microbial taxa and expanding our understanding of cyanobacterial diversity.

In recent years, the cyanobacterial family Oculatellaceae has garnered considerable attention, with numerous new genera and species being described, particularly from terrestrial and subaerial habitats. The type genus *Oculatella* is primarily soil-associated, though species have been identified in diverse environments such as caves, moist walls, and even anthropogenic substrates like clay pots. While some taxa occur in aquatic ecosystems, most members of Oculatellaceae exhibit a strong preference for drier environments (Akagha et al., 2023; Brito et al., 2022; Zammit, 2018).

Among the notable discoveries within this family are species such as *Oculatella subterranea* and *Oculatella kauaiensis*, both of which were isolated from cave ecosystems, as well as several taxa adapted to extreme soil conditions. The recently characterized genus *Egbenema* further exemplifies this adaptability, with species documented in Nigeria, Puerto Rico, and the Chihuahuan Desert—suggesting a broad ecological distribution (Akagha et al., 2023). However, despite its intriguing biogeography, research on *Egbenema* remains limited, and to date, no complete genome sequences have been published for any species within this genus.

Molecular studies have revealed that several Oculatellaceae species are morphologically cryptic yet genetically distinct, as evidenced by 16S rRNA gene sequencing. This exemplifies an ongoing challenge in cyanobacterial taxonomy: the rapid accumulation of genetic data far outpaces the formal description of taxa, leaving many so-called “dark taxa” undescribed (Li et al., 2020; Akagha et al., 2023).

In this study, we aimed to bridge these gaps by studying cyanobacteria from terrestrial habitats in the Northern Western Ghats. One of the enrichments was morphologically and molecularly distinct from known taxa. Using a combination of morphological analysis, 16S rRNA sequencing, and whole-genome sequencing, we identified this organism as a new species within the genus *Egbenema*. This study thus introduces a new species of *Egbenema* from the Indian subcontinent and marks a significant milestone in the genomic exploration of this genus. Characterizing such novel species further opens up the discovery of new bioactive compounds and genetic traits that could contribute to advancements in biotechnology and ecology.

## Methods

### Strain Isolation

The biological sample was obtained in June 2019 by gently scraping the bark surface of a Rain tree located in Nerul, Navi Mumbai, Maharashtra, India (19.0338° N, 73.0196° E) (Fig. 1). The isolation procedure began with an enrichment phase in a liquid BG11 medium supplemented with 1.5 g L⁻¹ NaNO₃ (Stanier et al., 1971). Upon the observation of visible microbial growth, filamentous structures were meticulously transferred to fresh liquid media, and a subset was streaked onto solid BG11 agar plates (1.5% w/v agar, Himedia). The process of transferring small amounts of biomass to new BG11 plates, followed by reintroduction into liquid medium, was repeated as necessary. Throughout these steps, all cultures were maintained under a 12-hour light/dark cycle with an illumination intensity of 20 μmol photons m⁻² s⁻¹ at a constant temperature of 28°C. Microscopic examinations were performed before and after each transfer, for both liquid and solid cultures, to ensure the effectiveness of the isolation process. Once isolated, the unicyanobacterial species was continuously cultured under the same controlled conditions.

**Fig 1:**
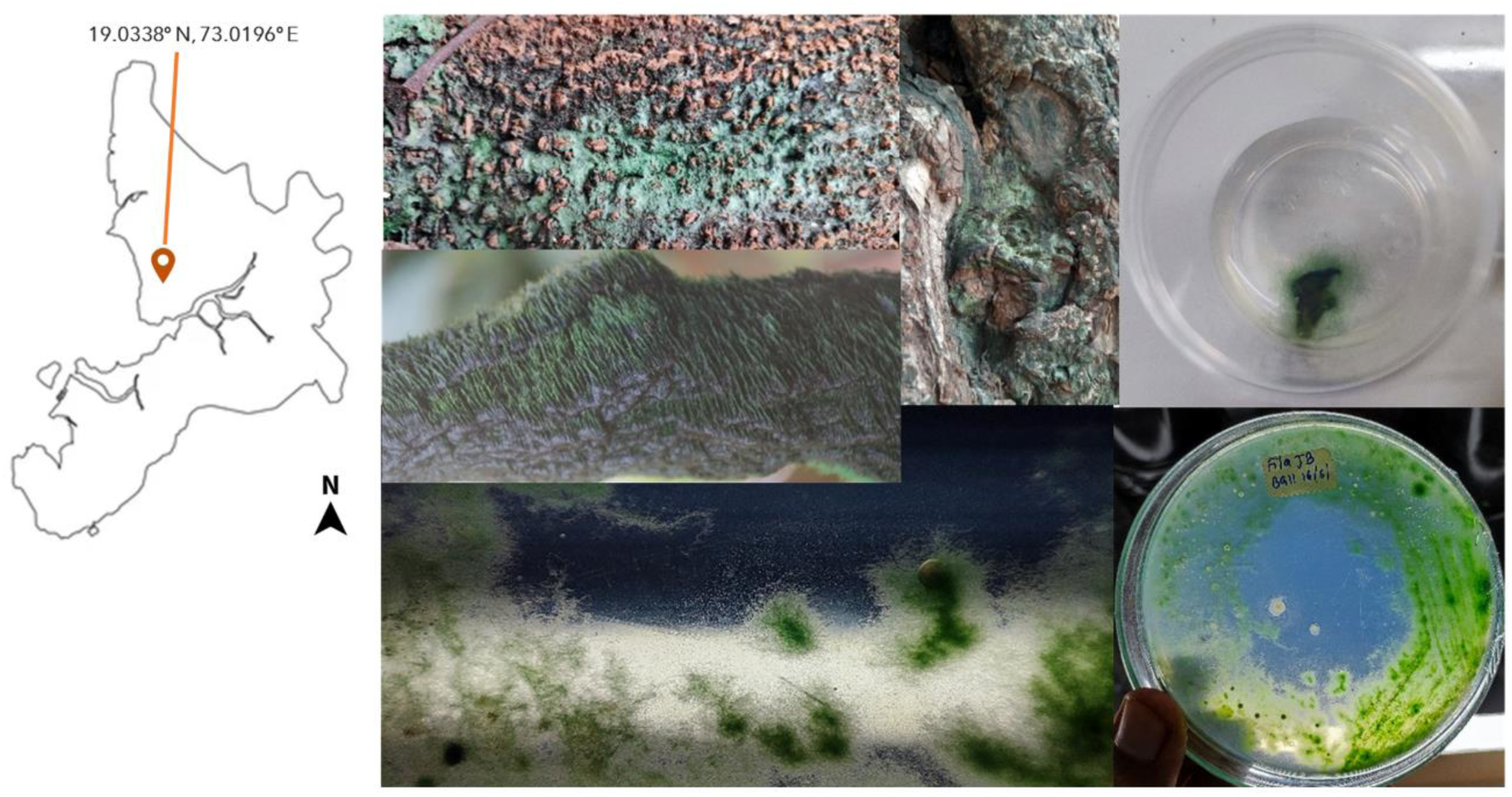
Geolocation and isolation of *Egbenema bharatensis*: from the surface of rain trees, enrichment and colony appearance clockwise.

### Morphological assessment

Cell morphology was examined using an Olympus Phase Contrast INVI Microscope, with images captured via its camera module and processed using ScopeImage 9.0 software. For a more detailed study, strain morphology was further analyzed on a Zeiss AxioImager Z1 light microscope equipped with an EC Plan-Neofluar 100×/1.3 N.A. oil immersion objective and DIC optics. High-resolution images were recorded with an AxioCam ERc 5s camera and managed using Zen 3 Pro software. Parameters evaluated included cell shape, terminal cell characteristics, dimensions (measured from 100 cells), reproductive features, presence of sheaths, branching patterns, and cell granulation.

### 16S rRNA molecular characterization

Genomic DNA was extracted from the cyanobacterial sample using the DNeasy® Plant Pro Kit (Qiagen, Hilden, Germany) in accordance with the manufacturer’s protocol. DNA concentration was determined with a DS-11+ spectrophotometer (DeNovix, Wilmington, USA). Partial 16S rDNA was amplified using cyanobacteria-specific primers (CYA106F, CYA781Ra, and CYA781Rb) in a 50 μl PCR reaction. This reaction comprised 25 μl of GoTaq® Green master mix (2X; Promega, Madison, USA), 5 μl of 10 µM forward primer (CYA106F), 2.5 μl each of 10 µM reverse primers (CYA781Ra and CYA781Rb), 1 μl of template DNA (112.538 ng/μl), and 14 μl of nuclease-free water. PCR cycling conditions were as follows: an initial denaturation at 94°C for 5 minutes; 35 cycles of 94°C for 1 minute, 60°C for 1 minute, and 72°C for 1 minute; and a final extension at 72°C for 10 minutes. Successful amplification was confirmed by agarose gel electrophoresis, with visualization carried out using a Gel Doc XR+ system (Bio-Rad, California, USA). The PCR product was then purified using the GENECLEAN® Turbo for PCR Kit (MP Biomedicals, California, USA), and its concentration was re-assessed with the DS-11+ spectrophotometer. Sequencing of the purified product was conducted using the Big Dye Terminator v3.1 cycle sequencing kit (Applied Biosystems, Foster City, USA) on a Genetic Analyzer 3130xl (Applied Biosystems, Foster City, USA), utilizing the three aforementioned primers. Sequence data were analyzed with FinchTv 1.4.0 (Geospiza Inc., USA), and the resulting contigs were manually assembled to generate a partial 16S rDNA sequence of 654 bp. This sequence was subsequently submitted to NCBI BLASTn for taxonomic identification.

### Sample Preparation

Genomic DNA was isolated from pure cultures using the SPINeasy® DNA Kit for Tissue & Bacteria, following an optimized protocol for high molecular weight DNA extraction. The quality and quantity of the extracted DNA were evaluated using a Qubit fluorometer and NanoDrop spectrophotometer, respectively. The integrity of the DNA was further confirmed by agarose gel electrophoresis, ensuring its suitability for downstream sequencing applications.

### Library Preparation

Whole genome sequencing libraries were prepared using the Rapid Barcoding Kit (SQK-RBK114.96) from Oxford Nanopore Technologies using the following protocol. 1 microgram of DNA from each sample was used as input. The DNA was first fragmented enzymatically using the transposase provided in the kit. Barcoding adapters were then ligated to the fragmented DNA to allow for sample differentiation during sequencing. The barcoded samples were pooled equimolarly to achieve uniform coverage across samples. The pooled library was loaded onto a MinION Flow Cell (FLO-MIN114) with an R10.4 nanopore configuration. Prior to sequencing, the flow cell quality was checked using the MinKNOW software, ensuring sufficient active pores were available. Sequencing was performed on a GridION MK1 device, with real-time basecalling enabled through the MinKNOW software. The run was monitored periodically, and sequencing was allowed to proceed until sufficient depth of coverage was achieved.

### Genome Assembly

The de novo genome assembly and annotation were performed using Oxford Nanopore Technologies’ Bacterial Assembly and Annotation Workflow (wf-bacterial-genomes), incorporating all necessary steps for bacterial genome analysis. Sequencing reads were first filtered to remove those shorter than 1000 bp or with a Q score below 10, ensuring high-quality input data. Assembly was conducted using the Flye 2.9.3 assembler, optimized for long-read data, followed by polishing with Medaka 1.12 to correct sequencing errors and enhance accuracy. Genome coverage and depth were assessed using Bamtools to confirm the completeness and reliability of the assembly. This comprehensive workflow ensures accurate and robust genome assembly.

### Genome Characterization

The genome sequence obtained was analysed using resources available at Bacterial and Viral Bioinformatics Resource Center (BV-BRC)(https://www.bv-brc.org/), Department of Energy Systems Biology Knowledgebase (KBase)(https://www.kbase.us/), and Rapid Annotations using Subsystems Technology (RAST) servers(https://rast.nmpdr.org/rast.cgi). Briefly the workflow of the analysis was as follows; Ensuring that the genome assembly is of high quality, with minimal gaps and errors. Contamination checks were performed using CheckM-v1.0.18 lineage workflow. Genome with good quality scores was annotated to identify genes and functional elements using Prokka - v1.14.5 and re-annotated using RASTtk - v1.073. The Genome was visualized and plotted using Circos. Furthermore, to identify metabolically relevant genes, pathway analysis was performed using KOALA (https://www.kegg.jp/blastkoala/) and to identify and analyze biosynthetic gene clusters (BGCs) within the novel genome, we utilized the antiSMASH 7.1.0 tool(https://antismash.secondarymetabolites.org/#!/start), which facilitates comprehensive detection and annotation of secondary metabolite biosynthesis pathways by comparing genomic sequences against a curated database of known BGCs (Blin et al., 2023). This approach allowed us to predict potential natural products and explore the biosynthetic capabilities of the organism. Additionally, identification and classification of toxin-antitoxin (TA) systems in the cyanobacteria genome were performed using TADB 3.0 (http://bioinfo-mml.sjtu.edu.cn/TADB3/)(Guan et al., 2024), to gain insights into survival strategies and stress responses.

### Comparative genome analysis

The genome was subjected to comprehensive genome analysis (CGA) to pinpoint its closest relatives using the Mash and MinHash tools. The genomes identified through CGA were then analyzed with FastANI v0.1.3 to measure genetic relatedness among prokaryotes, with ANI serving as a key metric for defining species boundaries. To support the FastANI findings, genomic similarity was also evaluated using the Genome-to-Genome Distance Calculator, which computes digital DNA-DNA Hybridization (dDDH) values.

Taxonomic classification of the novel genome was performed using the Genome Taxonomy Database Toolkit (GTDB-Tk). This tool compares genomes against a curated reference database through ANI and multiple sequence alignments (Chaumeil et al., 2022; Parks et al., 2022) ensuring a standardized taxonomic placement and accurate determination of phylogenetic relationships.

To confirm the isolate’s genus and species, the full-length 16S rRNA sequence retrieved from RAST annotation was queried against the NCBI nr/nt database using BLASTn. Additionally, the 16S-23S rRNA and ITS sequences were analyzed for their RNA secondary structure.

### ITS structure analysis

The ITS D1–D1′, Box–B, V2, and V3 helices were identified based on conserved basal clamp regions and their typical positions within the ITS (Akagha et al., 2023). Genome sequences from closely related *Egbenema* species were obtained from the same study to maintain consistency. Multiple sequence alignment, performed with the MUSCLE program known for its high accuracy, was used to precisely locate the ITS region in *Egbenema bharatensis* and related species. This alignment enabled a detailed comparison of the ITS regions across the species. The secondary structure of the ITS region was then predicted using the RNA fold form program, accessed through UNAFold web server, an online platform dedicated to Nucleic Acid Folding and hybridization (Zuker, 2003).

## Results

### Morphological Description

*Egbenema bharatensis* J. M. Bhagat, gen. et sp. nov. (Fig. 2a - g)

**Fig. 2:**
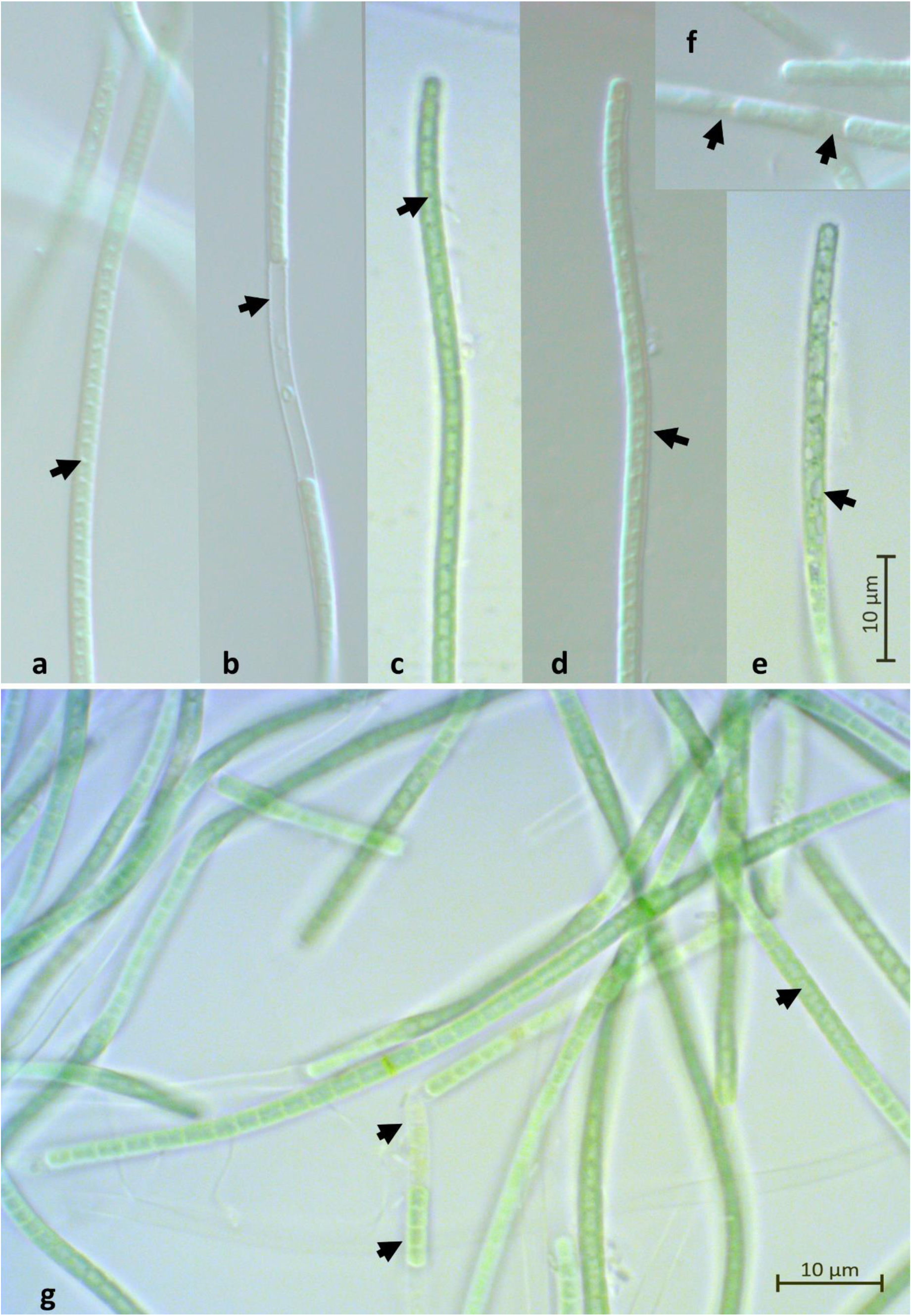
Morphological characteristic of *Egbenema bharatensis:* Cylindrical trichomes, not attenuated at the ends, show slight or no constrictions at cross-walls, with rounded apical cells that are longer or equivalent in length to interior cells (a, d). The sheath is thin, colorless, and occasionally exceeds the trichome (b, d). Cellular content reveals a visible peripheral chromatoplasm and central pale nucleoplasm, often containing granules at cross-walls, with aerotopes absent (c, e, g). Trichome fragmentation and subsequent disintegration lead to the release of hormogonia, a primary mode of reproduction (b, f, g).

HOLOTYPE: The holotype is a permanent fixed sample deposited in the Blatter Herbarium, St. Xavier’s College, Mumbai, Maharashtra, India, with the identifier Microbial Mats No. FJBBOT/2019.

TYPE STRAIN: The type strain is designated as FJBBOT/2019, and it is maintained in the culture collection of the Department of Botany, K. J. Somaiya College of Science and Commerce, Mumbai, Maharashtra, India.

TYPE LOCALITY: This species was isolated from a microbial mat on the barks of rain tree (*Samanea saman*) in Navi Mumbai, Maharashtra, India, by Jayesh M. Bhagat in June 2019.

ETYMOLOGY: The genus name “*Egbenema*” is derived from the location Egbe, Nigeria as described by Akagha et al,. 2023. The species name “*bharatensis*” refers to Bharat, the traditional name for India, signifying the geographical origin of this new cyanobacterial species.

### Description of New species

Colonies dark green to blue-green during the log phase, later turning yellow in the late stationary phase, flat mat, not penetrating agar. The filaments in the colony appear arranged in a loose, tangled mass rather than distinct fascicles or tufts. Filaments are green to blue-green, straight or curved. Sheath is colorless, thin (Fig. 2d), and not distinctly visible, except when exceeding the trichome (Fig.2b).

Trichomes are cylindrical in shape, not attenuated at the ends, slightly or not constricted at cross-walls (Fig. 2a, d), and immotile. False branching is absent. Cells are isodiametric or longer than wide, measuring 1.8–2.38 µm (average 2.07 µm) wide and 1.63–3.13 µm (average 2.25 µm) long. The apical cell is rounded, without a calyptra measuring average 2.48 µm long, longer or equivalent to interior cells. Heterocyst and Akinetes are absent.

Cell content is divided into a visible peripheral chromatoplasm and a central pale nucleoplasm, often containing granules at the cross walls and lacking aerotopes (Fig. 2c, e, g). Reproduction occurs by necridic cells (occasionally) and through trichome fragmentation, followed by disintegration, releasing hormogonia (Fig. 2b, f, g).

### 16S rRNA Molecular Identification

The partial 16S rRNA analysis showed 97.55% similarity to *Egbenema gypsiphylum*. To confirm the identity of the genera and species since the whole genome sequence for this cyanobacteria was not available in the database, we sequenced our strain to whole genome sequencing. (Supplementary file; Table S2 for BLAST result).

### Genome Analysis

A total of 2.55 GB of sequencing data was obtained, with an N50 of approximately 6 kb and a mean Q score of 15. The genome assembly, performed using the ONT assembly workflow, resulted in an assembled genome with the largest contig measuring ∼7.5 Mb and a coverage depth consistently exceeding 200X across the genome. Post-assembly quality control included contaminant screening to ensure data reliability. Evaluation of the *Egbenema bharatensis* genome using **CheckM** (v1.018) demonstrated a high-quality assembly with 98.94% completeness and 2.44% contamination, confirming its suitability for downstream applications.

### Gene content

The draft, near-complete genome spans approximately 7,506,658 bp and exhibits a G+C content of 50.34%. Initial annotation carried out using Prokka (v1.14.5) identified 6,839 coding sequences (CDSs) and 13,847 features. When reannotated with RASTtk, the number of CDSs increased to 6,947. In contrast, the RAST pipeline identified 8,200 CDSs and 8,253 features. Despite these differences, all three annotation tools consistently detected 53 RNAs, including the full set of tRNAs commonly observed in cyanobacterial genomes.

The RAST pipeline identified 1,111 genes (14%) in subsystems and 7,089 genes (86%) outside subsystems, totaling 8,200 genes. Of these, 5,065 (61.7%) were classified as hypothetical proteins. The PATRIC database annotation similarly recognized 8,200 coding sequences (CDSs) and 9,240 features, aligning with the RAST-based structural annotations. Minor discrepancies between annotations likely result from differences in pipeline versions or reference databases.

Among the functionally annotated proteins, 1,073 carried Enzyme Commission (EC) numbers, 924 possessed Gene Ontology (GO) terms, and 817 were linked to KEGG pathways (data available upon request). Overall, 51.3% of the CDSs were predicted to encode proteins with discernible biological functions, while the remaining 48.7% were designated as hypothetical proteins. The genome also harbors 10 genes involved in nitrogen fixation (*nifB, nifS, nifN, nifH, nifE, nifT, nifX, nifU, nifZ,* and *nifW*).

The genome of *Egbenema bharantensis* contains total 591 genes for CRISPR, 7 confirmed CRISPRs CDS, 283 CRISPR repeats with 276 CRISPR spacers. Gene annotation using PATRIC database resulted in maximum number of annotated genes and functional assignments as compared to Prokka and RAST due to their inherent customisation in the pipelines used.

Furthermore, AntiSMASH analysis revealed five putative biosynthetic gene clusters in total, one of which exhibited similarity to an anachlain biosynthetic gene cluster, as determined by the ClusterBlast feature within AntiSMASH (Supplementary Fig. S1).

### Functional Analysis

Functional annotation using clusters of orthologous genes (COGs) revealed that a notable portion of the identified genes remain poorly characterized. Among those annotated, most were linked to metabolism, followed by genes involved in cellular processes and signaling, information storage and processing, and multifunctional categories (Supplementary Fig. S5). The selection of eggNOG mapper for functional annotation was based on its well-established ability to rapidly and accurately annotate new sequences, whether genes or proteins. This tool leverages pre-computed orthology assignments from the comprehensive eggNOG database, providing reliable functional predictions grounded in evolutionary relationships. Compared to simple homology-based searches, eggNOG mapper offers a more robust, context-aware framework, making it particularly well-suited for annotating genomes from poorly characterized or novel microorganisms (Cantalapiedra et al., 2021; Huerta-Cepas et al., 2019).

The genomic analysis of this terrestrial cyanobacterium revealed a complex network of toxin-antitoxin (TA) systems, comprising 46 identified genes—93.5% of which are Type II and 6.5% Type VII. These systems include well-known families such as VapC/VapB, HicA/HicB, and RelE/RelB, all recognized for their roles in stress response and cell cycle regulation. Additionally, genes associated with plasmid stability (parD, RHH, CopG), metal ion transport (mntA), and potential biotechnological applications (PIN domain, MazF) were identified. Beyond these, 19 rarely occurring genes were detected that may possess unique functions. This diverse collection of TA systems suggests that the cyanobacterium has a robust set of tools for adapting to various environmental stresses and competing within microbial communities (Supplementary Fig. S6, Table S1).

The functional analysis of annotated genes reveals a diverse metabolic repertoire with significant biotechnological potential. Notably, the presence of pathways for cobalamin (vitamin B12) synthesis, heme biosynthesis, and nitrogen fixation hints towards the bacterium’s capacity for producing valuable compounds. The detection of respiratory complex I and pyruvate metabolism pathways further suggest potential for bioenergy generation and biofuel production. Moreover, the identification of genes related to cysteine synthesis and stress response including heavy metal tolerance correlates with the organism’s habitat resilience and adaptability, positioning it as a promising candidate for industrial applications. By harnessing these metabolic capabilities, this cyanobacterium could contribute to sustainable solutions in sectors such as agriculture, healthcare, and energy production.

### Comparative genome analysis

The closest neighbour genome to *Egbenema bharatensis* as determined by two methods Mash and MinHash and through Species tree(49 core, universal genes defined by COG gene families) revealed it to be closest to the available genome of [Leptolyngbya_sp._JSC_1_GCF_000733415.1] refer Fig. 3. FastANI analysis revealed below threshold values of 77.78% ANI values between the novel genome and *Leptolyngbya_sp*._JSC_1_GCF_000733415.1, thus not supporting its placement within the same genus boundary. This finding was further corroborated by the digital DNA-DNA Hybridization (dDDH) results from the Genome-to-Genome Distance Calculator, the *Egbenema bharatensis* genome is not conspecific or consubspecific with any of the reference genomes. Logistic regression models calculate a **0% probability** of the query genome being the same species or subspecies as any of the reference genomes. The significant G+C content differences (>1%) and consistently low DDH values (12.5% to 26%) corroborate the conclusion that the genome likely represents a novel Genera and species.

**Fig. 3:**
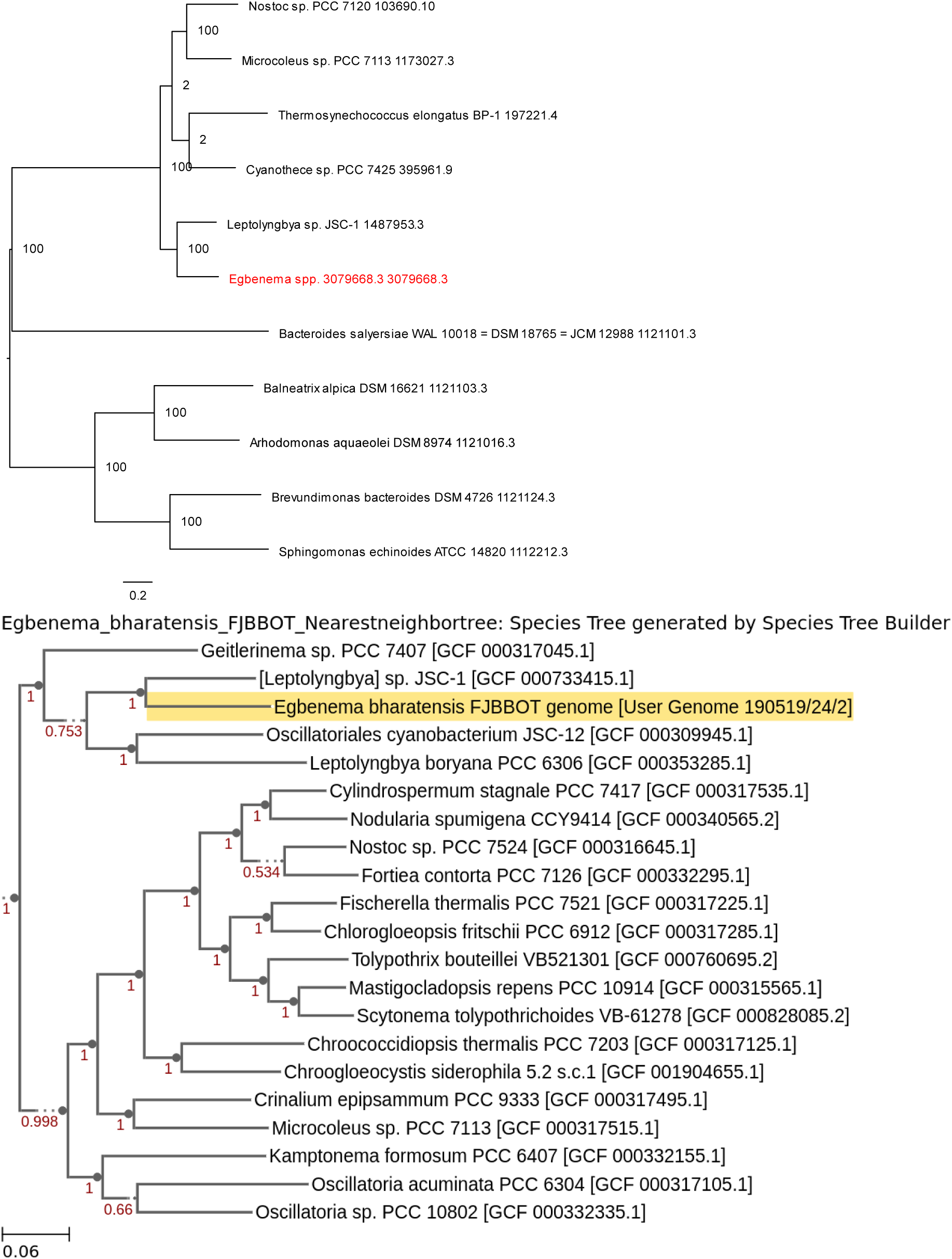
Phylogenomic tree of *Egbenema bharatensis* using a) Mash/MinHash and b)Codon tree methods

The Taxonomic classification using GTDB-Tk placed the genome within d Bacteria; p Cyanobacteriota; c Cyanobacteriia; o Elainellales; f Elainellaceae; g CCP2; s CCP2 sp003003615, confirming its phylogenetic consistency and novelty.

BLASTn analysis of the full-length 16S rRNA sequence identified *Egbenema gypsiphilum* strain WHSA1-4-NP1A as the most similar genus, with sequence similarity of 97.64%. Refer table 3.

### ITS secondary structure analysis

The secondary structure of the ITS region of *Egbenema bharatensis* includes all the conserved helices, such as D1-D1’, Box-B, and V3, as found in the genera *Egbenema*, and is compared with other species (Fig. 4). The D1-D1’ helix is structurally similar to *Egbenema sp. CY40 KY421773* exhibiting the characteristic unmatched cytosine residue opposite the basal 3′ unilateral bulge and the enlarged terminal loop and unmatched A at 16^th^ position. Box-B helix is structurally similar to *Egbenema sp. CHAB* but showed marked differences in both sequence as well as structure with other species. The V3 helix has two loops, terminal and basal region, which is identical to *Egbenema sp. CY40 KY421773*.

**Fig. 4.**
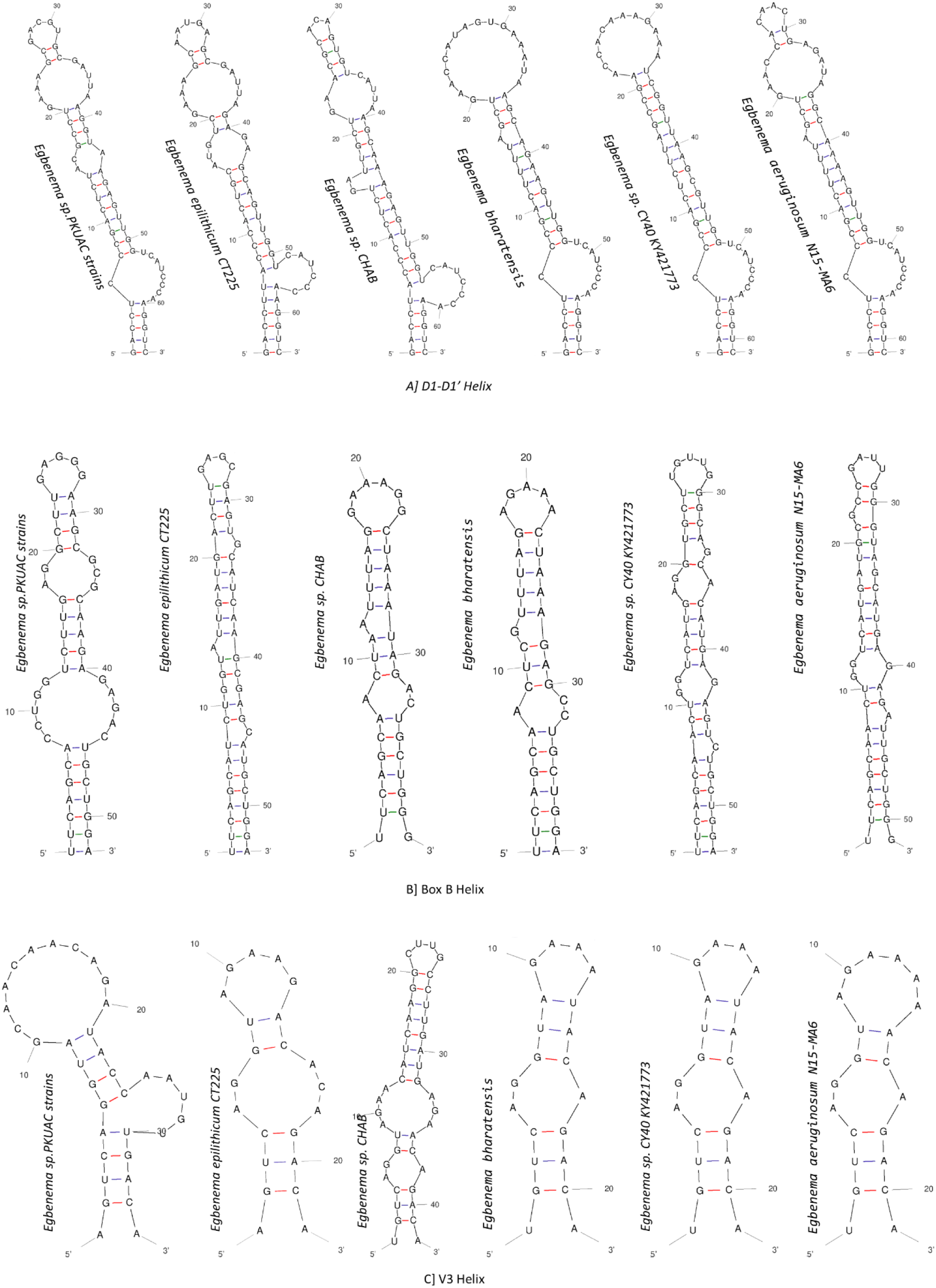
ITS secondary structure comparison; A] D1-D1’ helix, B] Box-B helix, C] V3 helix

## Discussion

The discovery of a novel species within the genus *Egbenema* emphazises the vast and underexplored microbial diversity of the Northern Western Ghats, particularly in terrestrial habitats, which remain underrepresented in biodiversity studies compared to aquatic ecosystems. Cyanobacterial diversity presents unique challenges for classification due to their asexual reproduction, morphological convergence, and the limitations of traditional morphology-based methods (Dvořák et al., 2015; Johansen & Casamatta, 2005; Ward et al., 2021). Advances in genomics and molecular techniques—such as whole-genome sequencing and metabarcoding—have greatly refined our ability to delimit species, revealing cryptic genera and species that once went unnoticed (Li et al., 2020). There is now a clear need to standardize cyanobacterial nomenclature using genome-based phylogeny, which in turn calls for high-quality genome sequences to improve classification (Dextro et al., 2021). Our study shows how these modern tools can help uncover new taxa and deepen our understanding of phylogenetic relationships in biodiversity hotspots like the Western Ghats.

The genus *Egbenema* is a striking example of ecological adaptability. Its members thrive in a range of environments: *Egbenema aeruginosum*, the type species, is well-adapted to Nigeria’s tropical forests; *Egbenema epilithicum* flourishes in the tropical monsoonal climate of Puerto Rico; and *Egbenema gypsiphilum* has been found in the arid gypsum landscapes of the Chihuahuan Desert (Akagha et al., 2023). This broad distribution speaks to the versatility of *Egbenema* species and their ability to survive under varied environmental conditions. The discovery of a new *Egbenema* species in habitats heavily impacted by human activity in the Northern Western Ghats not only aligns with previous sampling but also hints at the potential for further diversification in these largely unexplored areas.

A detailed morphological comparison of *Egbenema bharatensis* with other species (see Table 1) reveals distinct characteristics—such as the absence of false branching, differences in cell size, and the presence of necridia—that set this species apart within the family. Although these morphological clues suggested the possibility of a new species, they alone were not enough to conclusively identify it.

**Table 1:** Morphological comparison of different species of *Egbenema*.

| Characteristic | *Egbenema aeruginosum* | *Egbenema epilithicum* | *Egbenema gypsiphilum* | *Egbenema bharatensis* J. M. Bhagat, gen. et sp. nov. |
| --- | --- | --- | --- | --- |
| Thallus/Colony | Blue-green, flat mat, not penetrating agar. | Bright blue-green, flat mat, not penetrating agar. | Bright blue-green, floating mat. | Dark green to blue-green in log phase, yellow in stationary phase, flat mat, not penetrating agar. |
| Filaments | Straight, single trichome per sheath, infrequent false branching, 2.2–2.8 µm wide. | Straight and coiled, single trichome per sheath, frequent false branching, 1.8–3.0 µm wide. | Straight and coiled, frequent false branching, 2.2–2.8 µm wide. | Straight or curved, arranged in a loose, tangled mass, not in distinct fascicles, no branching, 1.8–2.38 µm wide. |
| Sheath | Narrow, clear, tightly adherent, occasionally exceeding end cells, facultatively present. | Narrow, clear, tightly adherent, occasionally exceeding end cells, facultatively present. | Narrow, clear, tightly adherent, occasionally exceeding end cells, facultatively present. | Thin, colorless, not distinctly visible, occasionally exceeding trichomes, facultatively present. |
| Trichomes | Slightly constricted at crosswalls, 2.0–2.6 µm wide. | Slightly or not constricted at crosswalls, 1.6–3.0 µm wide. | Mostly constricted at crosswalls, unconstricted within sheath, 2.0–2.6 µm wide. | Cylindrical, not attenuated at ends, slightly or not constricted at crosswalls, immotile, 1.73–2.63 µm wide. |
| Cells | Bright blue-green, parietal thylakoids along outside walls, 1.2–2.8 µm long. | Blue-green, parietal thylakoids along outside walls, 1.6–4.0 µm long. | Blue-green, parietal thylakoids along outside walls, often with large granules, 1.8–3.6 µm long. | Blue-green, isodiametric or longer than wide, visible peripheral chromatoplasm, central pale nucleoplasm, 1.63–3.13 µm long. |
| Apical Cells | Slightly longer than interior cells, 1.8–3.2 µm long. | Longer or shorter than interior cells, 2.0–4.0 µm long. | Longer or shorter than interior cells, yellowish, 2.0–4.2 µm long. | Rounded, without calyptra. Longer than or equal to interior cells, 2.2- 3.25 $\mu$ m long |
| Granules/Inclusions | Lacking granules. | Without granules. | Frequently with a single large granule and reddish inclusions toward apices. | Granules present at cross walls, lacking aerotopes. |
| Reproduction | Necridia unobserved. | Necridia unobserved, hormogonia formed by fragmentation. | Rare necridia, hormogonia formed by fragmentation. | Rare necridic cells, via trichome fragmentation releasing hormogonia. |

Initial molecular characterization using cyanobacteria-specific 16S rRNA primers confirmed the isolate’s affiliation with the genus *Egbenema*. However, the noticeable morphological differences and a sequence identity below the 98% threshold (Edgar, 2018) highlighted the limitations of conventional methods in resolving species-level taxonomic ambiguities. This limitation prompted us to turn to whole-genome sequencing—a powerful approach for both microbial taxonomy and evolutionary studies.

Whole-genome sequencing provided the detailed resolution needed to tackle these challenges, offering clear insights into the isolate’s taxonomic placement and filling an important gap in the genomic data available for the genus *Egbenema*. Comparative genomic analyses were then carried out to rule out the possibility that the genome belonged to another genus. While the analysis initially placed the genome near *Leptolyngbya JSC1*, a cyanobacterial taxon from a different order, further validation using genomic similarity indices such as ANI and GGDC revealed a low genetic similarity between the two. This evidence reinforces the novelty of the isolate. Moreover, the nearest neighbor predicted by GTDB taxonomy was identified as *Ellainella*, a related genus within the family Oculatellaceae—a misclassification that further emphasizes the challenges posed by incomplete genomic databases. This situation clearly demonstrates the need for more comprehensive genomic data to accurately resolve taxonomic relationships within the family (Dextro et al., 2021; Lounici et al., 2024; Simonazzi et al., 2024).

Since comparative genomics strategies confirmed that the genome significantly doesn’t match with any known genomes or reference genome, we analysed further the full-length 16S rRNA sequence and ITS region, which provided critical insights into the isolate’s taxonomic identity. The full length 16S rRNA Blast results also pointed to the genus *Egbenema* while again failing to meet the species-level identity threshold. The ITS region in such cases proves critical for distinguishing species due to its higher variability compared to the 16S rRNA gene. Integrating phylogenomic data with ITS analysis has proven invaluable for refining taxonomic classifications and identifying newly described taxa (Strunecký et al., 2023). In our study, ITS secondary structure analysis uncovered distinctive features that set this isolate apart from other *Egbenema* species, further supporting its designation as a new species within the genus.

This study is significant in two major ways. First, it presents the first complete genome sequence for *Egbenema*, filling a crucial gap in the genomic resources available for this genus. Complete genomes are essential for understanding the evolutionary relationships, functional capabilities, and ecological roles of microorganisms (Chen et al., 2021; Dextro et al., 2023). Second, the discovery of a novel species from the Indian subcontinent emphasizes the value of exploring underexamined environments—particularly terrestrial habitats in biodiversity hotspots and areas influenced by human activity—to uncover hidden microbial diversity.

Our findings have broader implications for cyanobacterial taxonomy and phylogeny as well. The family *Oculatellaceae* has experienced extensive taxonomic revisions, with recent research revealing many cryptic species and underscoring the need for integrative, polyphasic approaches to classification (Akagha et al., 2023). This work illustrates the utility of relying on fixed sequence similarity thresholds for species identification as an indicator and demonstrates the benefits of combining morphological, molecular, and genomic data to achieve taxonomic resolution.

Moreover, the genomic data revealed several unique features in the new isolate, including a full complement of genes for nitrogen fixation and an unusually high number of toxin-antitoxin systems. These findings suggest that the novel *Egbenema* species may possess ecological or biotechnological potential, warranting further experimental studies to explore its functional attributes which are underway and would be published in subsequent communication.

## Conclusion

In conclusion, this work enriches our understanding of the genus *Egbenema*, introducing a novel species *Egbenema bharatensis* from the Indian subcontinent and expanding the genomic representation of this ecologically versatile genus. The findings reinforce the importance of integrating morphological, molecular, and genomic approaches to uncover and characterize microbial diversity. Furthermore, the study highlights the Northern Western Ghats as a critical region for microbial exploration, emphasizing the need for continued research into its diverse and underexplored habitats.

## Acknowledgments

The authors thank Dr. Deepti Gupta for primer recommendation and MTCC, Punjab for partial 16S sequencing and molecular identification of the cyanobacteria. We thank Dr. Khushboo Gandhi, Ms. Vaishali, ACTREC, Kharghar for assisting in imaging. We also thank Somaiya Vidyavihar University, Mumbai for providing the necessary infrastructural support for this research work.

## Consent for publication

All the authors have read the paper and given their approval for it to be published in this journal.

## Disclosure statement

No potential conflict of interest was reported by the authors.

## Funding

The study was not funded by any funding agencies including the public, commercial, or not-for-profit sectors.

## Data availability statement

This Whole Genome sequencing project has been deposited at NCBI Genome database under the accession id PRJNA1221537.

## Author Contribution

**JB** Conceptualization, Methodology, Software, Validation, Formal Analysis, Investigation, Resources, Data Curation, Writing - Original Draft, Writing - Review & Editing, Visualization, Supervision. **AK** Conceptualization, Methodology, Software, Validation, Formal Analysis, Investigation, Resources, Data Curation, Writing - Review & Editing. **DS** Methodology, Software **MP** Conceptualization Writing - Review & Editing, Supervision.

## Declaration of generative AI and AI-assisted technologies in the writing process

During the preparation of this work the author(s) used GPT-4o service in order to improve the clarity and language of the writing. After using this tool/service, the author(s) reviewed and edited the content as needed and take(s) full responsibility for the content of the published article.

## Supplementary File

**Fig. S1:**
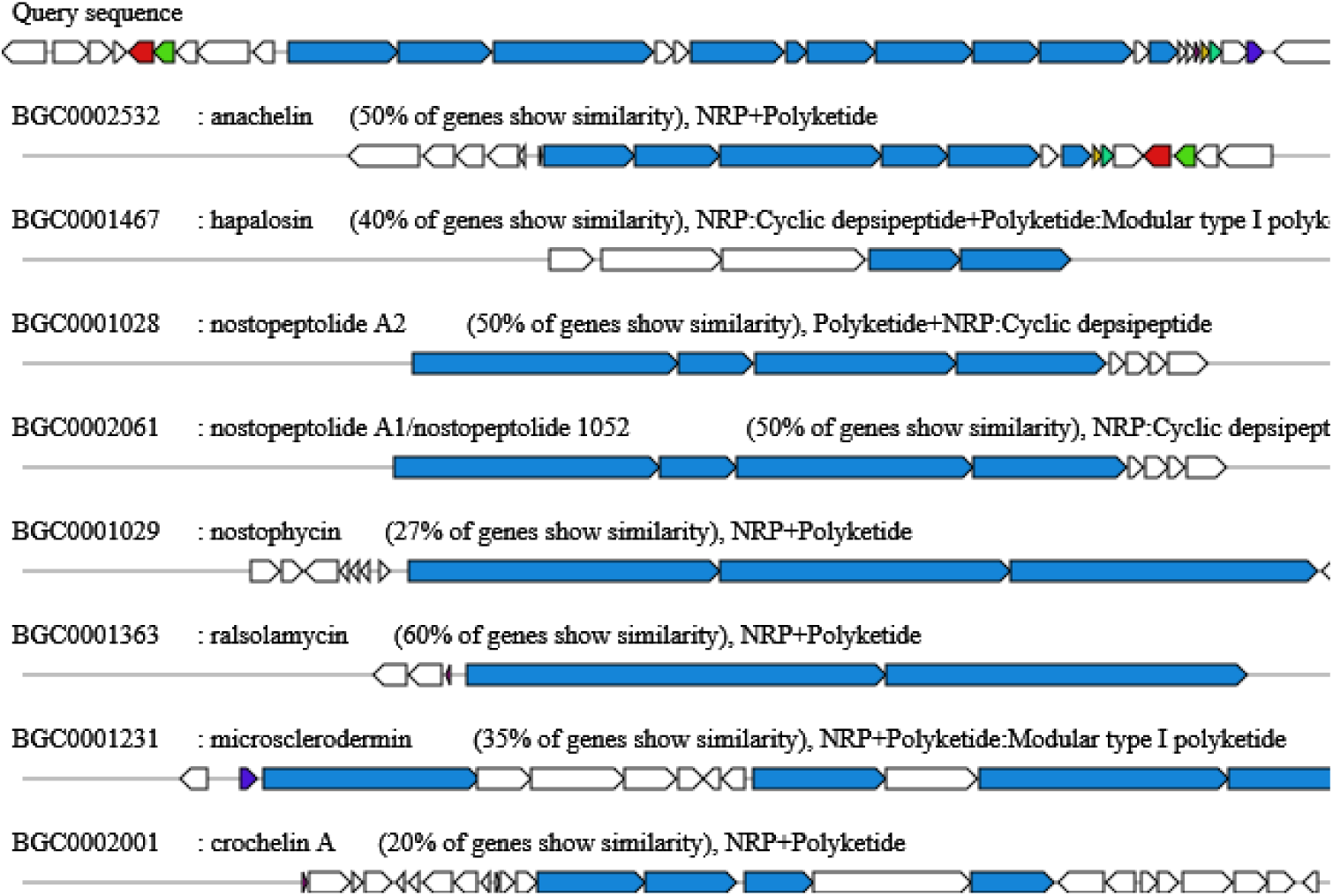
BGC gene cluster prediction results

**Fig. S2.**
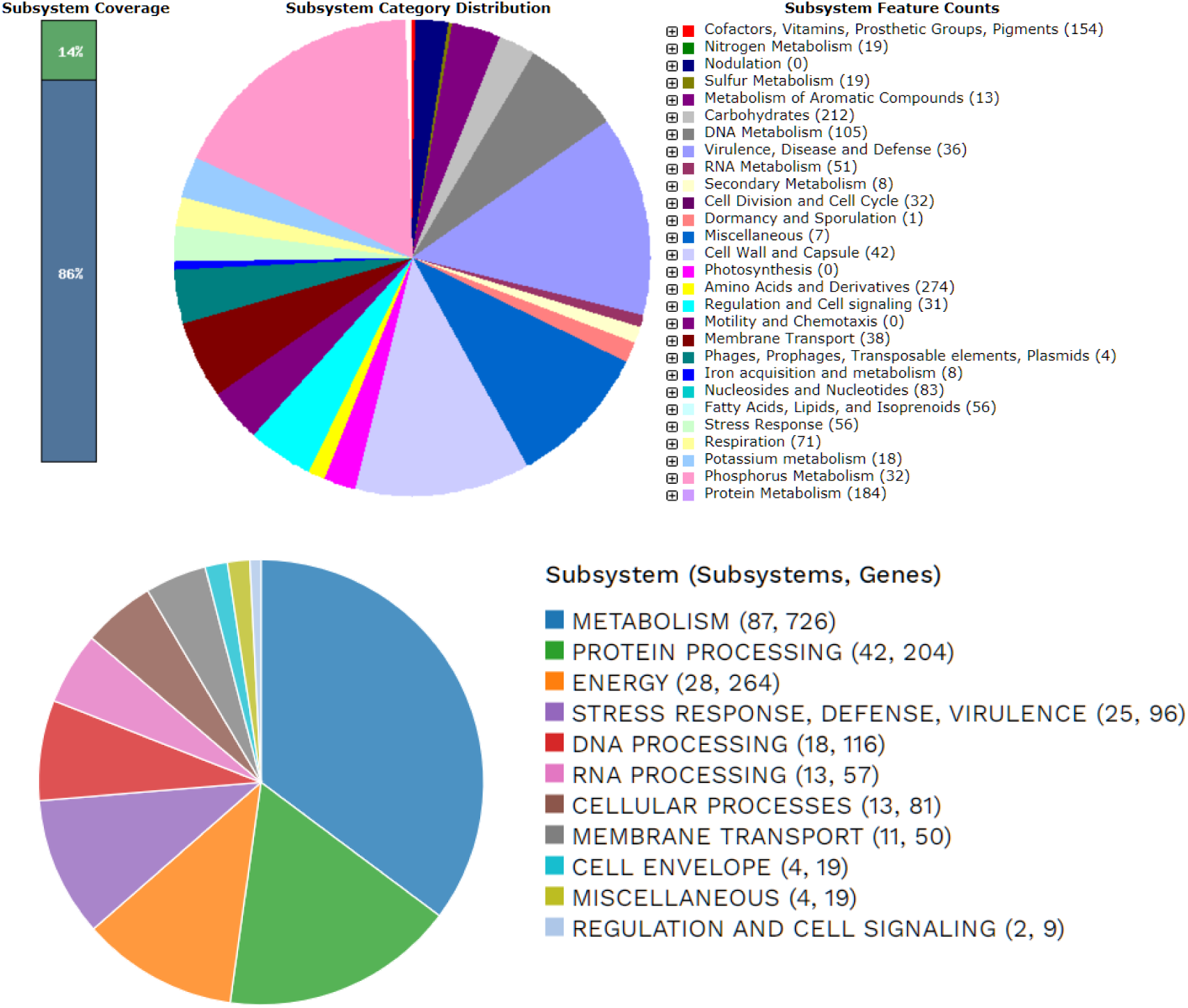
Subsystem distribution as annotated by RAST server and PATRIC annotation

**Fig. S3:**
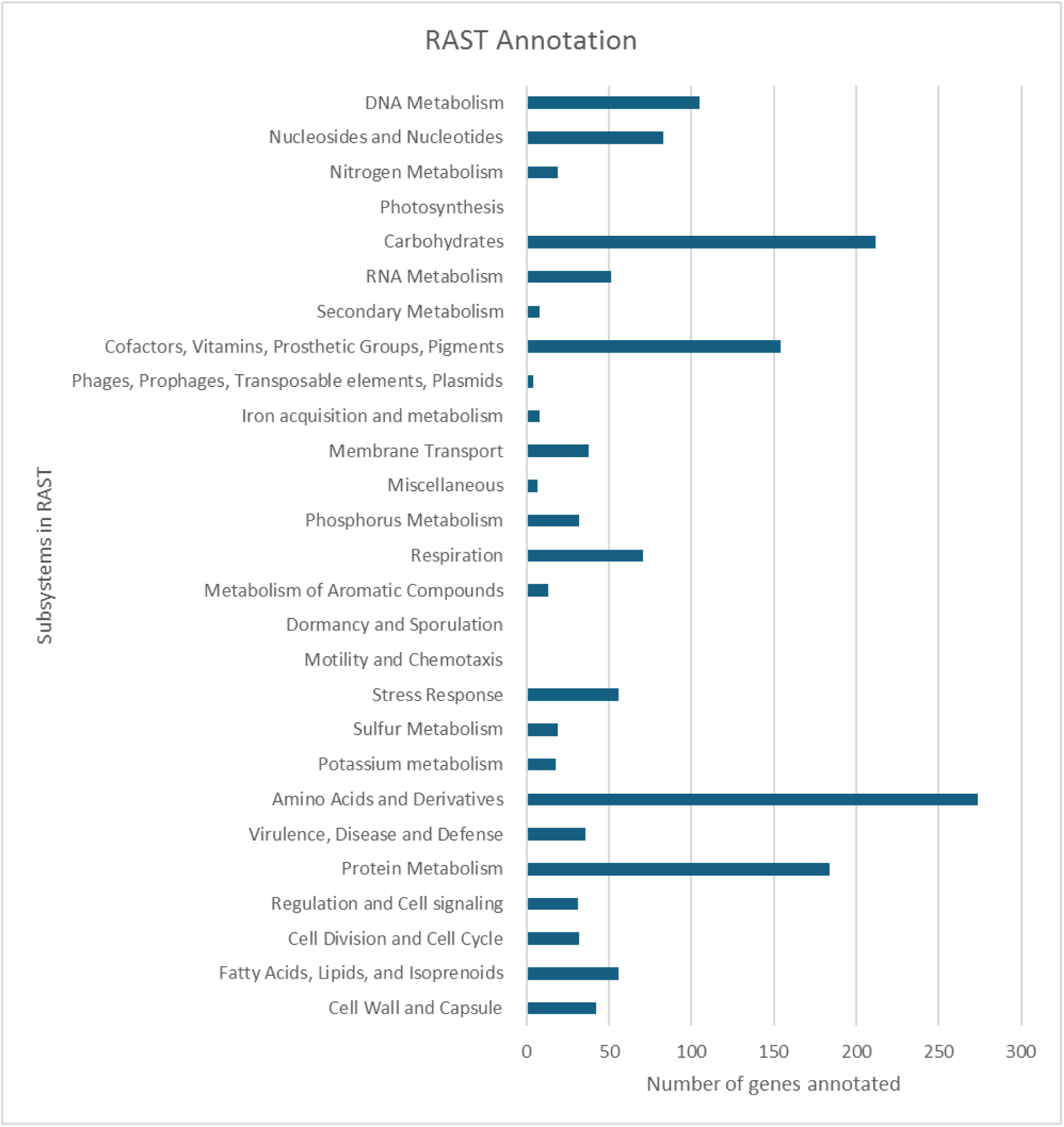
Distribution of genes annotated through RAST subsystem.

**Fig. S4:**
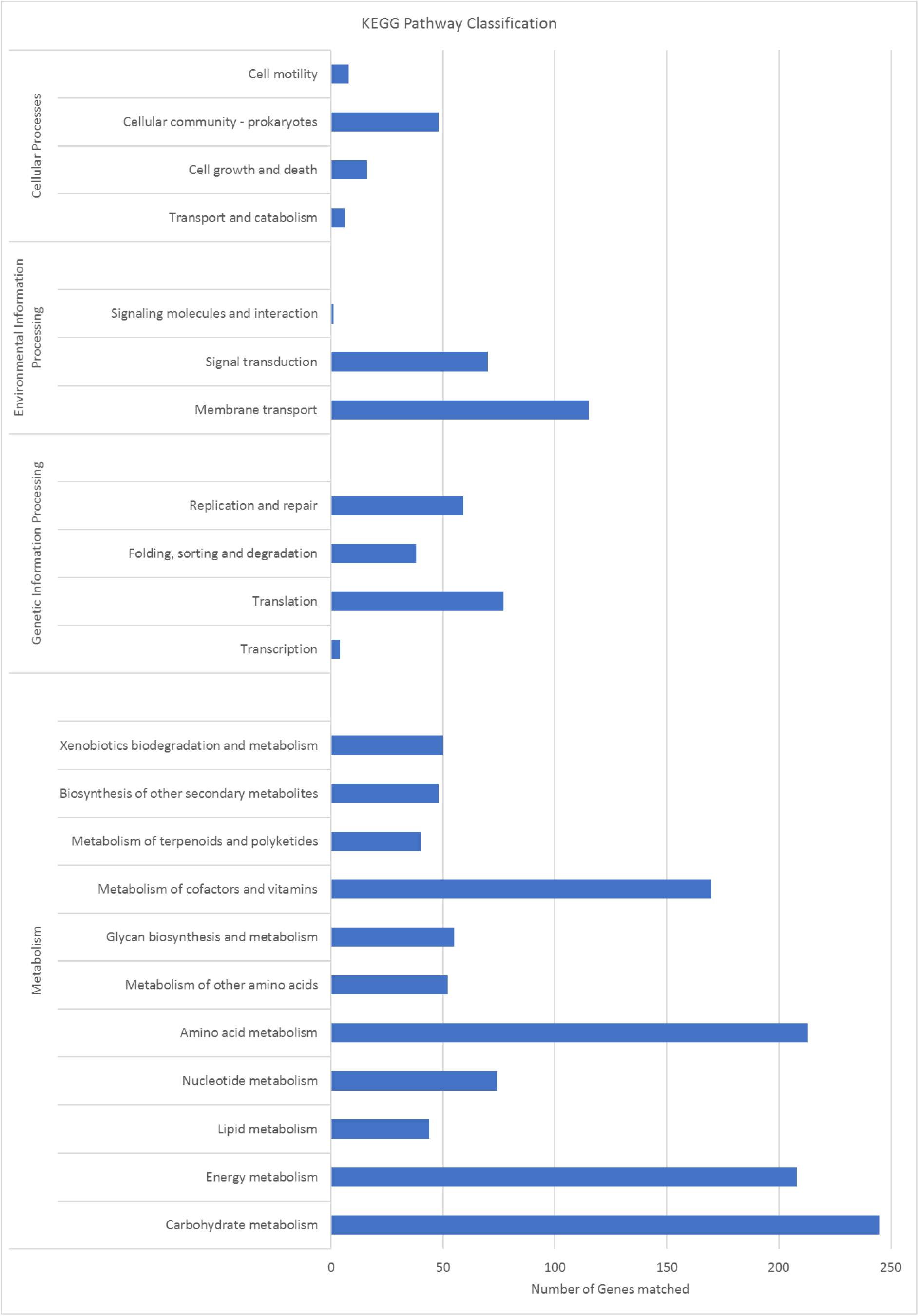
KEGG pathway enrichment; revealing the metabolic potential of *Egbenema bharatensis*

**Fig. S5:**
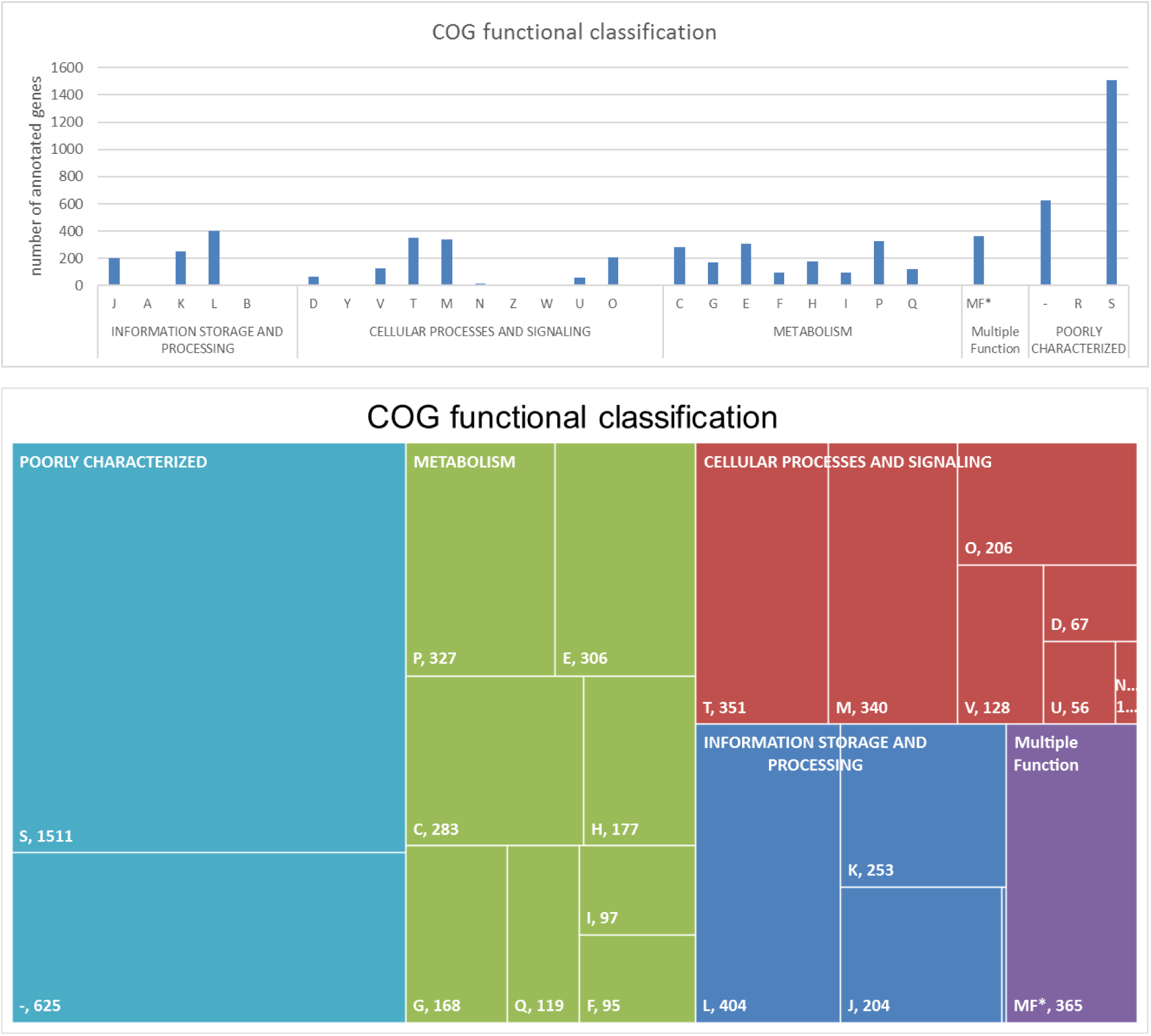
Functional classification of proteins annotated using eggNOG mapper.

**Fig. S6:**
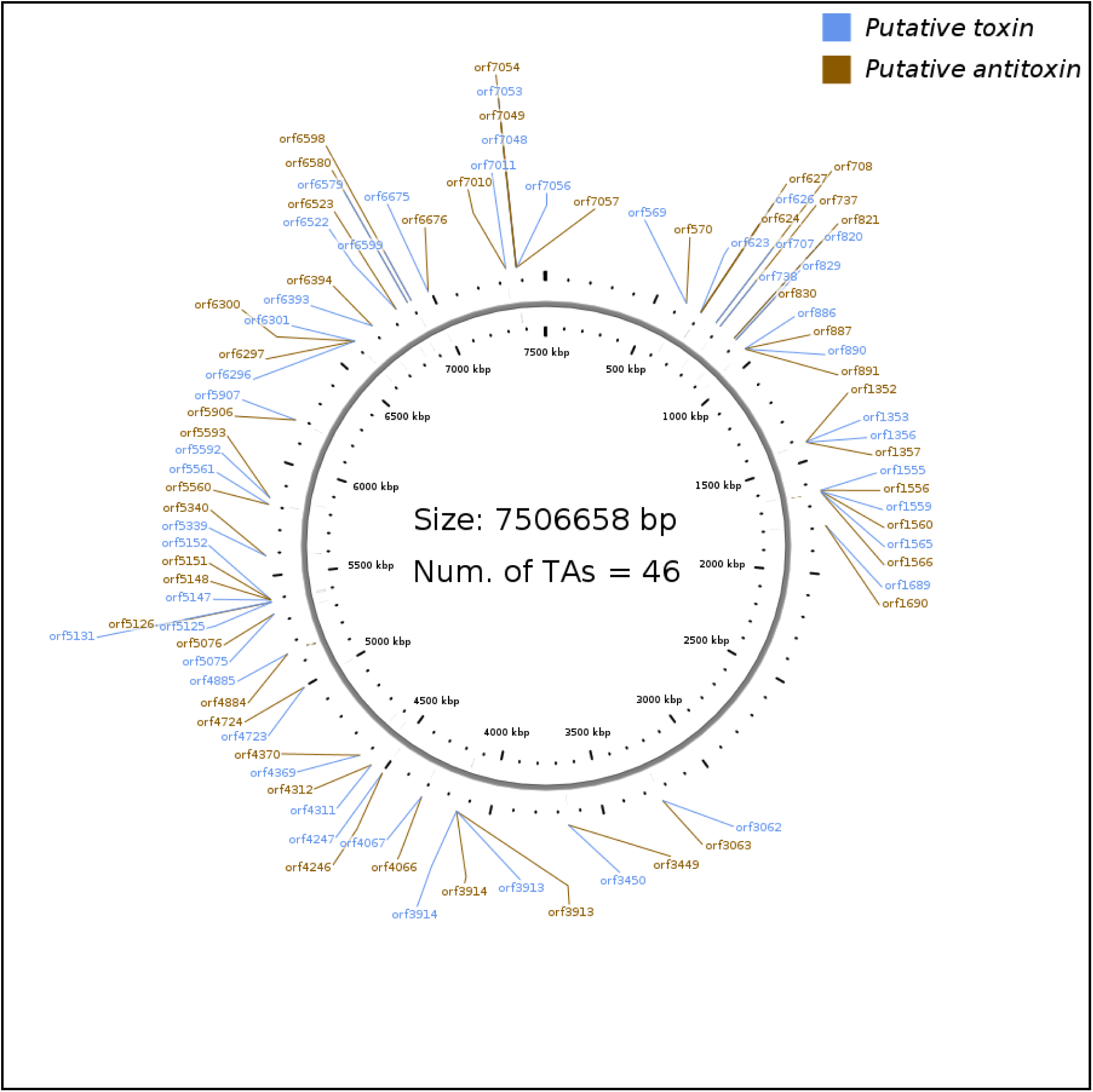
TA mapping with genome of *Egbenema bharatensis*

**Table S1:** Toxin Antitoxin gene occurrences in *Egbenema bharatensis*.

| Gene Type | Occurrences | Category Description |
| --- | --- | --- |
| vapC | 12 | Toxin component of TA systems; inhibits translation. |
| vapB | 10 | Antitoxin that neutralizes VapC toxin. |
| hicB | 7 | Antitoxin, often regulates HicA toxin. |
| hicA | 7 | Toxin affecting mRNA stability. |
| mntA | 4 | Metal ion transporter (Mn/Fe-specific). |
| HicA | 4 | Another annotation of hicA toxin. |
| PIN | 4 | Domain present in RNase-related toxins. |
| relE | 3 | Toxin; cleaves mRNA at ribosomes under stress. |
| mazE | 3 | Antitoxin neutralizing mazF toxin. |
| DUF3368 | 3 | Domain of unknown function in TA system components. |
| relB | 3 | Antitoxin paired with RelE toxin. |
| BrnT | 2 | Toxin related to cell cycle control. |
| CopG | 2 | Regulator in plasmid stability mechanisms. |
| RHH | 2 | Ribbon-helix-helix motif; DNA-binding domain. |
| mazF | 2 | Toxin inhibiting translation via mRNA cleavage. |
| parD | 2 | Antitoxin associated with plasmid stabilization. |
| UPF0175 | 2 | Protein family related to stress response. |
| hepT | 2 | Toxin-antitoxin gene pair regulator. |
| Others (19 genes) | 1 each | Rarely occurring genes with unique functions. MNTss, HEPN, vagC, higB, higA, VapC, Protein of unknown function (DUF2281), fip/doc, toxT, sezA, MarR, pezT, yoeB, parE, Uncharacterised protein family (UPF0175), AbrB/MazE, nucleic acid-binding protein, contains PIN domain,StbC |

**Table S2:**
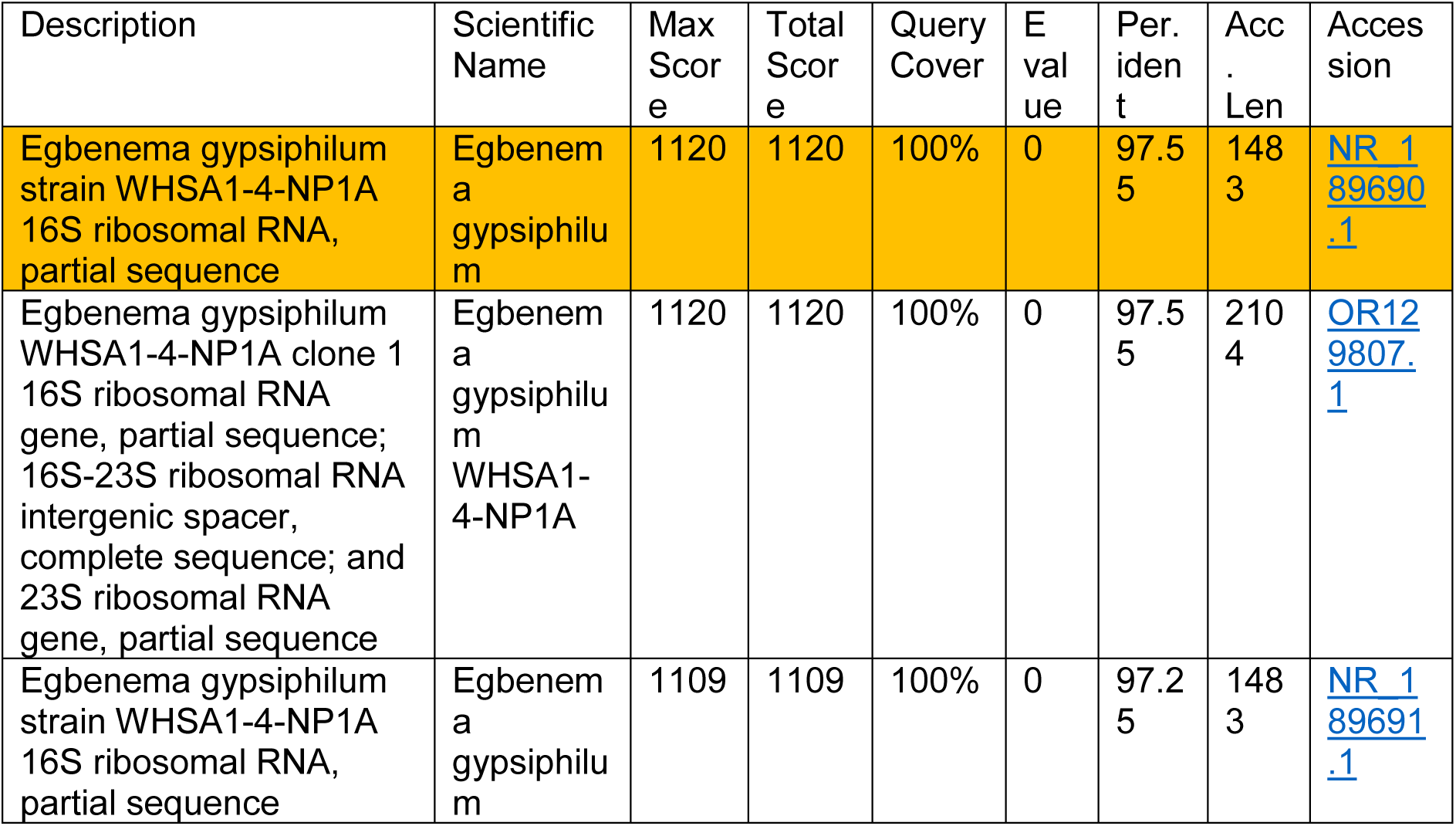
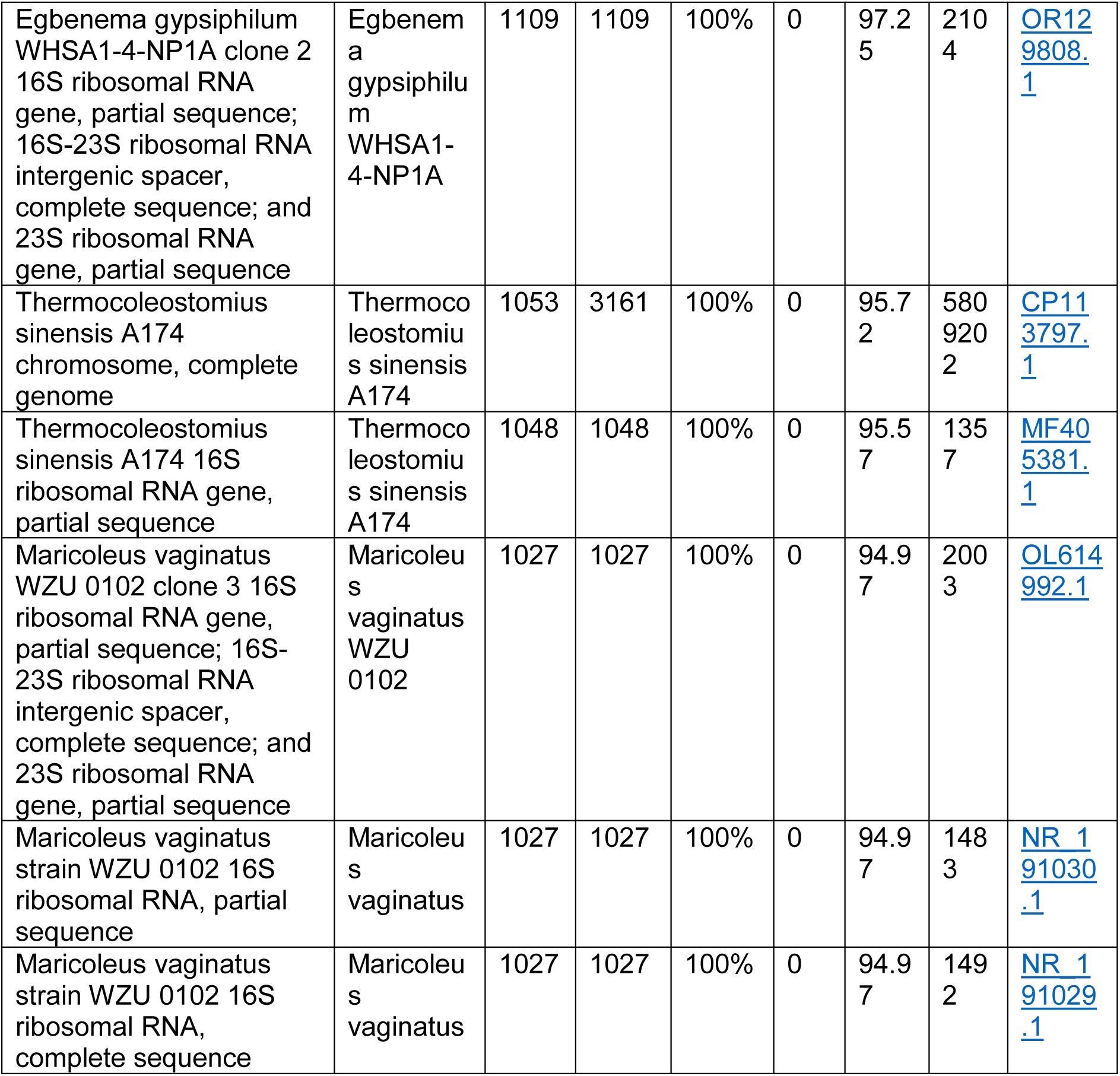
Blastn output description of *Egbenema bharatensis* partial 16S rRNA sequence.

| Description | Scientific Name | Max Score | Total Score | Query Cover | E value | Per. ident | Acc. Len | Accession |
| --- | --- | --- | --- | --- | --- | --- | --- | --- |
| Egbenema gypsiphilum strain WHSA1-4-NP1A 16S ribosomal RNA, partial sequence | Egbenema gypsiphilum | 1120 | 1120 | 100% | 0 | 97.55 | 1483 | NR_189690.1 |
| Egbenema gypsiphilum WHSA1-4-NP1A clone 1 16S ribosomal RNA gene, partial sequence; 16S-23S ribosomal RNA intergenic spacer, complete sequence; and 23S ribosomal RNA gene, partial sequence | Egbenema gypsiphilum WHSA1-4-NP1A | 1120 | 1120 | 100% | 0 | 97.55 | 2104 | OR129807.1 |
| Egbenema gypsiphilum strain WHSA1-4-NP1A 16S ribosomal RNA, partial sequence | Egbenema gypsiphilum | 1109 | 1109 | 100% | 0 | 97.25 | 1483 | NR_189691.1 |
| Egbenema gypsiphilum WHSA1-4-NP1A clone 2 16S ribosomal RNA gene, partial sequence; 16S-23S ribosomal RNA intergenic spacer, complete sequence; and 23S ribosomal RNA gene, partial sequence | Egbenema gypsiphilum WHSA1-4-NP1A | 1109 | 1109 | 100% | 0 | 97.25 | 2104 | OR129808.1 |
| Thermocoleostomius sinensis A174 chromosome, complete genome | Thermocoleostomius sinensis A174 | 1053 | 3161 | 100% | 0 | 95.72 | 5809202 | CP113797.1 |
| Thermocoleostomius sinensis A174 16S ribosomal RNA gene, partial sequence | Thermocoleostomius sinensis A174 | 1048 | 1048 | 100% | 0 | 95.57 | 1357 | MF405381.1 |
| Maricoleus vaginatus WZU 0102 clone 3 16S ribosomal RNA gene, partial sequence; 16S-23S ribosomal RNA intergenic spacer, complete sequence; and 23S ribosomal RNA gene, partial sequence | Maricoleus vaginatus WZU 0102 | 1027 | 1027 | 100% | 0 | 94.97 | 2003 | OL614992.1 |
| Maricoleus vaginatus strain WZU 0102 16S ribosomal RNA, partial sequence | Maricoleus vaginatus | 1027 | 1027 | 100% | 0 | 94.97 | 1483 | NR_191030.1 |
| Maricoleus vaginatus strain WZU 0102 16S ribosomal RNA, complete sequence | Maricoleus vaginatus | 1027 | 1027 | 100% | 0 | 94.97 | 1492 | NR_191029.1 |

**Table S3:** Blastn output description of *Egbenema bharatensis* complete 16S rRNA sequence.

| Description | Scientific Name | Max Score | Total Score | Query Cover | E value | Per. ident | Acc. Len | Accession |
| --- | --- | --- | --- | --- | --- | --- | --- | --- |
| Egbenema gypsiphilum strain WHSA1-4-NP1A 16S ribosomal RNA, partial sequence | Egbenema gypsiphilum | 2540 | 2540 | 98% | 0 | 97.64 | 1483 | NR_189691.1 |
| Egbenema gypsiphilum strain WHSA1-4-NP1A 16S ribosomal RNA, partial sequence | Egbenema gypsiphilum | 2540 | 2540 | 98% | 0 | 97.64 | 1483 | NR_189690.1 |
| Maricoleus vaginatus strain WZU 0102 16S ribosomal RNA, complete sequence | Maricoleus vaginatus | 2300 | 2300 | 99% | 0 | 94.63 | 1492 | NR_191029.1 |
| Maricoleus vaginatus strain WZU 0102 16S ribosomal RNA, partial sequence | Maricoleus vaginatus | 2296 | 2296 | 98% | 0 | 94.62 | 1484 | NR_191031.1 |
| Maricoleus vaginatus strain WZU 0102 16S ribosomal RNA, partial sequence | Maricoleus vaginatus | 2296 | 2296 | 98% | 0 | 94.62 | 1483 | NR_191030.1 |
| Maricoleus vaginatus strain WZU 0102 16S ribosomal RNA, partial sequence | Maricoleus vaginatus | 2289 | 2289 | 98% | 0 | 94.56 | 1483 | NR_191028.1 |
| Siamcapillus rubidus strain NUACC13 16S ribosomal RNA, partial sequence | Siamcapillus rubidus | 2242 | 2242 | 98% | 0 | 94.18 | 1539 | NR_186891.1 |
| Shahulinema minutum strain SH01 16S ribosomal RNA, partial sequence | Shahulinema minutum | 2239 | 2239 | 99% | 0 | 93.89 | 1486 | NR_191044.1 |
| Albertania obscura strain BACA0713 16S ribosomal RNA, partial sequence | Albertania obscura | 2239 | 2239 | 97% | 0 | 94.29 | 1465 | NR_191036.1 |

## Notes

### Competing Interest Statement

The authors have declared no competing interest.

